# Career Navigator: An online platform to streamline professional development and career education for graduate bioscientists

**DOI:** 10.1101/2024.02.22.580689

**Authors:** Rachel M. Rudlaff, Utsarga Adhikary, Candrika D. Khairani, Daniel S. Emmans, Johanna L. Gutlerner, Ronald J. Heustis

## Abstract

Graduate professional development is a highly dynamic enterprise that prepares graduate students for personal and career success in a variety of fields, including the biosciences. National policies, funding awards, and institutional programs have generated myriad tools and services for graduate bioscience students, including new learning resources, events, connections to prospective employers, and opportunities to strengthen academic and professional portfolios. These interventions are welcome and have done much to enhance graduate bioscience training, but they may also be overwhelming for trainees. To streamline professional development and career education information for the bioscience graduate students at our institution, we tested a model where we built a centralized web portal of career development resources. Here we present our strategy and best practices for website design. We show data that students preferred a centralized online portal over other forms of resource communication; that programming, paired communication and environmental factors (e.g. remote learning and work as in the COVID-19 pandemic) combined to increase sustained engagement with the site; and that harnessing website analytics is an effective way to measure site utilization and generate insights on programming and resource development. This data, in turn, fits into broader priorities to evaluate interventions in graduate bioscience education.

## INTRODUCTION

Graduate bioscience professional development supports the preparation of masters and doctoral trainees for the variety of jobs they will pursue after completing their graduate degrees. Across institutions, this can encompass career counseling, individual development planning, job search coaching, and support for managing transitions. Over the past decade, there has been a much-needed focus on and invigoration of graduate bioscience professional development. In 2012, the Biomedical Research Workforce Working Group Report commissioned by the National Institutes of Health (NIH) [1] coincided with and supported an emphasis by the NIH on the now-widely used Individual Development Planning for graduate bioscience trainees [2]. This report also laid a foundation for a series of funding opportunities offered through the NIH for Broadening Experiences for Scientific Training (BEST), designed to spur innovation in career exploration and career preparation for graduate bioscience trainees. Evidence of the breadth and impact of these funding awards are now emerging; for example, among the 17 BEST recipient institutions the most common interventions introduced were single-day workshops, internships, and self-assessments for trainees [3]. These awards not only catalyzed innovations at recipient institutions, but also triggered activity across the full national landscape of graduate biomedical training programs, as other institutions sought to improve biomedical professional advising through funding from other mechanisms (e.g., from institutional sources). Furthermore, systemic change was driven by the largest of the NIH agencies, the National Institute of General Medical Sciences (NIGMS). Indeed, informed by a recognition that doctoral bioscience graduates now pursue a broad range of careers outside the traditional PhD to postdoc to PI pipeline, the NIGMS significantly revamped the criteria for renewing and retaining competitive training grants (T32s) by placing substantially greater emphasis on graduate professional development [4].

This awareness and resource development has ushered an emergence of myriad new resources and tools to guide graduate students through traditional aspects of career preparation. These include resources and tools pertaining to career exploration, experiential learning (including internships), networking, finding jobs and applying for employment in various sectors and functions. Career preparation has also expanded beyond these roles, to now frequently integrating and promoting trainee wellness and mental health; providing training on diversity, inclusion and belonging toward promoting proper academic and workplace interactions; and supporting trainees considering their own identity-driven needs and values as part of the career exploration and job search process (e.g., culturally aware job searching). This breadth of training is exemplified by offerings from the NIH’s Office of Intramural Training and Education [5].

However, this proliferation of materials is only effective if students know about them and can access them when needed. Emails are often used to advertise new content, but students often miss these notices, or do not recall them when they are looking for a resource or service. In contrast, organizing content to be continuously available and easily searchable for graduate students is likely to maximize student professional development. A key component of this organizational framework includes highlighting how these pieces of information relate to each other. Online platforms offer a pragmatic solution to accomplish this, as long as they have robust offerings and are designed to guide the user to access the information most important to their unique needs. Indeed, online platforms can facilitate that by collating relevant content available both on-campus and externally; online platforms not only offer ease of accessibility but can also offer a wide variety of opportunities in one place. To achieve the highest impact and utility, these sites must be actively maintained to stay up to date; must provide opportunities for human engagement (e.g., with staff or professionals); and must be open to feedback from all users.

Approaches that simultaneously tackle graduate student career preparation and mental health may be both feasible and warranted. Graduate student mental health and well-being are related to career preparation: negative feelings about career prospects have been closely linked to mental health issues in graduate students [6]. Recently, online platforms have been proposed as a solution towards connecting graduate students to mental health resources, but the effectiveness of such platforms will depend on both student engagement and the processes that keep the platforms current and curated [7]. The importance of embedding well-being considerations into the culture of training, highlighting the importance of well-being for and to mentors, and streamlining access to resources to promote student well-being have also been highlighted [8].

All of these recommendations are aligned with best practices on Individual Development Plans (IDPs) for graduate students. IDPs offer a mechanism to incorporate self-reflection and active planning for professional development and well-being; are a conduit for training on and discussions among mentors and trainees; and can benefit from an informed and strategic integration of this wealth of resources. IDPs may also provide additional benefits for minoritized students by improving mentor-mentor interactions, helping these students build additional connections that promote their professional growth and overcome imposter syndrome [9]. Ultimately, our goal is to streamline professional development and career education communication so that it can be most *efficiently* used by trainees, both for their direct benefit and also to address faculty concerns that investment in professional development activities is at the expense of academic and research development and productivity.

To best address the career and professional development needs of the students at our institution - specifically through easy and equitable access to resources for professional development and mental health that could benefit IDPs and mentor-mentee discussions - we set out to ask several questions:

(1.) Did our students see a need for a centralized career education portal? If so, did we have data to support this?
(2.) Could we define a web-based model to systematically harness this type of information from various schools at our institution and external sources, and keep it continually updated? In doing so, what best practices of website design could we implement? Could we integrate a taxonomy to organize content, particularly resources, to systematically organize the site?
(3.) How could we drive continued and repeated use of such a site?
(4.) How would we track and evaluate such engagement?

Addressing these needs for our students and answering these research questions culminated in our building of the Career Navigator, a one-stop service center connecting trainees with tools, resources, events, news and providers related to career skills and professional development. In this paper, we highlight data from students at our private, research-intensive (R1) institution - with a student body split between a main campus and medical campus - that supports the need for such a site. We share the design approach for building the site and discuss how we utilized events and communication channels to drive its continued and repeated use. We also discuss how we use analytics to understand the impact of our interventions. Through developing this platform, its underlying taxonomy to categorize resources, and a communication strategy to promote engagement with the Career Navigator, we demonstrate that a centralized graduate bioscience career planning platform can be used as an organized repository of materials to support graduate student career development.

## RESULTS

### Pre-launch student perceptions on value

We had access to standard doctoral program evaluation data which measured student perceptions of many aspects of their programs. One survey, 2-4 months before the site’s launch in August 2018, asked doctoral students to “*Please rate how effective each of the following methods is or would be to you, as a means of making you aware of resources concerning fellowships, career exploration events, internships, and job opportunities*’’ by ranking five modes of communication on a 5-pt Likert scale from *not at all effective* to *extremely effective*. The results from our population show that among students - in aggregate across all 14 life sciences doctoral programs and all years (e.g., second year [G2], fourth year [G4], etc.) - there was strong interest in having a single website that could house all career and professional development information (**Figure 1A**). 51.7% of responses highlighted that a centralized website would be very effective (VE) or extremely effective (EE), while 44.7% said that weekly emails from their departments or programs would be VE+EE. There was lower enthusiasm for a weekly digest from a central administrative office (VE+EE = 37.1%), separate emails for each announcement from their departments or programs (VE+EE=35.7%) or separate emails for each announcement from a central administrative office (VE+EE = 24.6%). We did not have access to similar or other relevant data from our master’s students. This was consistent with our anecdotal data that students had begun to “tune out” email messages and reminders given the frequency of messages, which were often redundantly sent at the school, program and affinity group levels. This data was consistent across all student years, including early-stage (G2 or G3) and students at later stages (**Figure 1B-E**).

**Figure 1.**
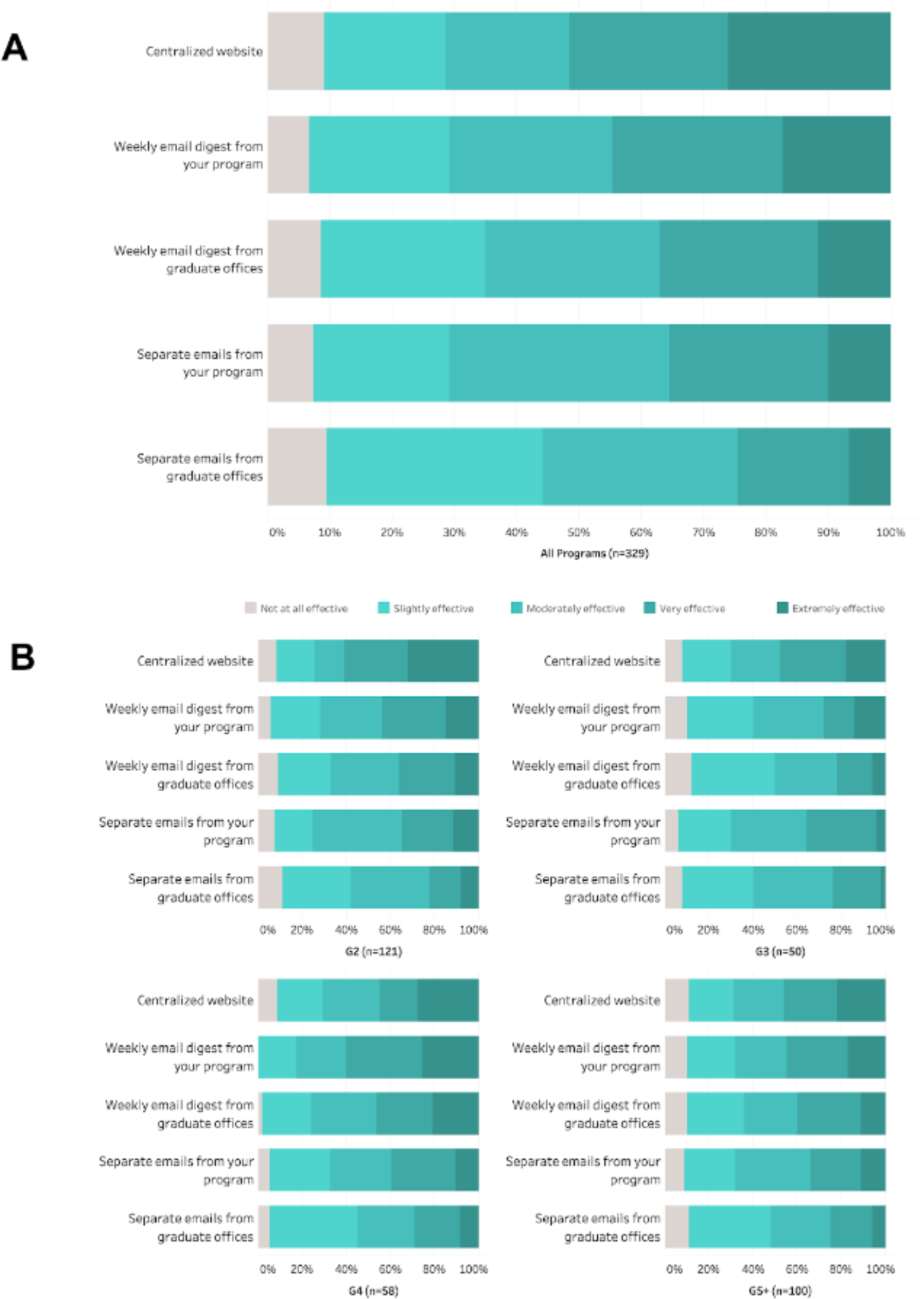
Students reported a preference for a centralized website housing all career education related information, as opposed to digest emails and regular emails for each opportunity. G2+ students (second year doctoral students and later stages) were surveyed during the period April 2018 to June 2018; at that time, >1000 students received the survey. **(A)** 329 students completed the entire survey which was the criteria for inclusion in the analysis. Pooling responses across all 14 of our life sciences doctoral programs shows that students reported a preference for a centralized website where all material relevant to career education was centralized. For comparison, we focused on responses as very effective (VE) or extremely effective (EE). We found that VE+EE was 51.7% of responses for a central website, 44.7% for weekly digests from their department or program, 37.1% for a weekly digest from central administration, 35.6% for separate emails from their department or program for each notice, and finally 24.7% for separate emails from central administrative office for each notice. **(B)** This preference was true across all levels of our graduate student population. The same results were observed with the cross-sectional analysis of students in their G2 year (second year doctoral), G3, G4 and G5+ (fifth year doctoral and beyond). Figures were generated using Tableau.

### Design, build, maintenance and driving engagement for the site

In developing the Career Navigator, we identified four channels for information to which we could assign notices stemming from within and outside our university. These channels became the sub-pages to our site, an application of “chunking” where similar information is grouped to define website architecture. As such, we proposed pages for (a) resources, (b) events, (c) job postings, and (d) announcements.

The “resources” page of our site was designed to house information meant to be “permanent” (i.e., not time-sensitive and enduringly relevant): it links users to tools (e.g., video libraries, websites, simulations) that do not have an expiration date (either existentially or in their relevance to users), as opposed to events, job postings, and (most) announcements which are time-specific.

The “events” page captures upcoming events, hosted within our university or from external organizations. Originally, all these events were hosted live, but since March 2020 - at the start of the remote learning and work due to the COVID-19 pandemic - our events have all been virtual (until September 2022 when we began offering both in-person events again along with virtual events). These have also been predominantly synchronous events.

Our “job postings” page is our space for compiling advertisements for short- and long-term employment opportunities, including postdoctoral positions. Importantly, we catalog postdoctoral opportunities not only in the sciences and scientific research, but in broader areas for these transitional stages. We include those designed for learning about and transitioning to careers in discipline-based education research (DBER); teaching and curriculum development; career advising and professional development; and, science policy and advocacy, among others. We list the search engines known for searching jobs relevant to graduate bioscience students (annotating them with their sectors of relevance). Since launch, we have noted the need for a place to post experiential learning opportunities (e.g., short-term internships) and we currently post these transient experiential learning opportunities to our “announcements” page.

“Announcements” is an all-encompassing page for content that is time-limited in relevance; for example, notices about approaching fellowship deadlines, on-campus jobs, internship application deadlines, volunteer opportunities, and the launch of new student organizations and other initiatives. This page is not for new resources, events, or job postings (as defined previously). New resources, events and job postings are relayed on the previously described relevant pages and have been highlighted via a bi-weekly newsletter (discussed below).

The Career Navigator organizes all these types of information onto a single website, including information from the various schools within our university as well as information emanating from external sources (**Figure 2**). In addition to the site, the Career Navigator has two additional pages: a “Home” page and “Contact Us” (the latter discussed below as part of maintenance). Interested users can access this publicly available site at https://careernavigator.gradeducation.hms.harvard.edu/ [10].

**Figure 2.**
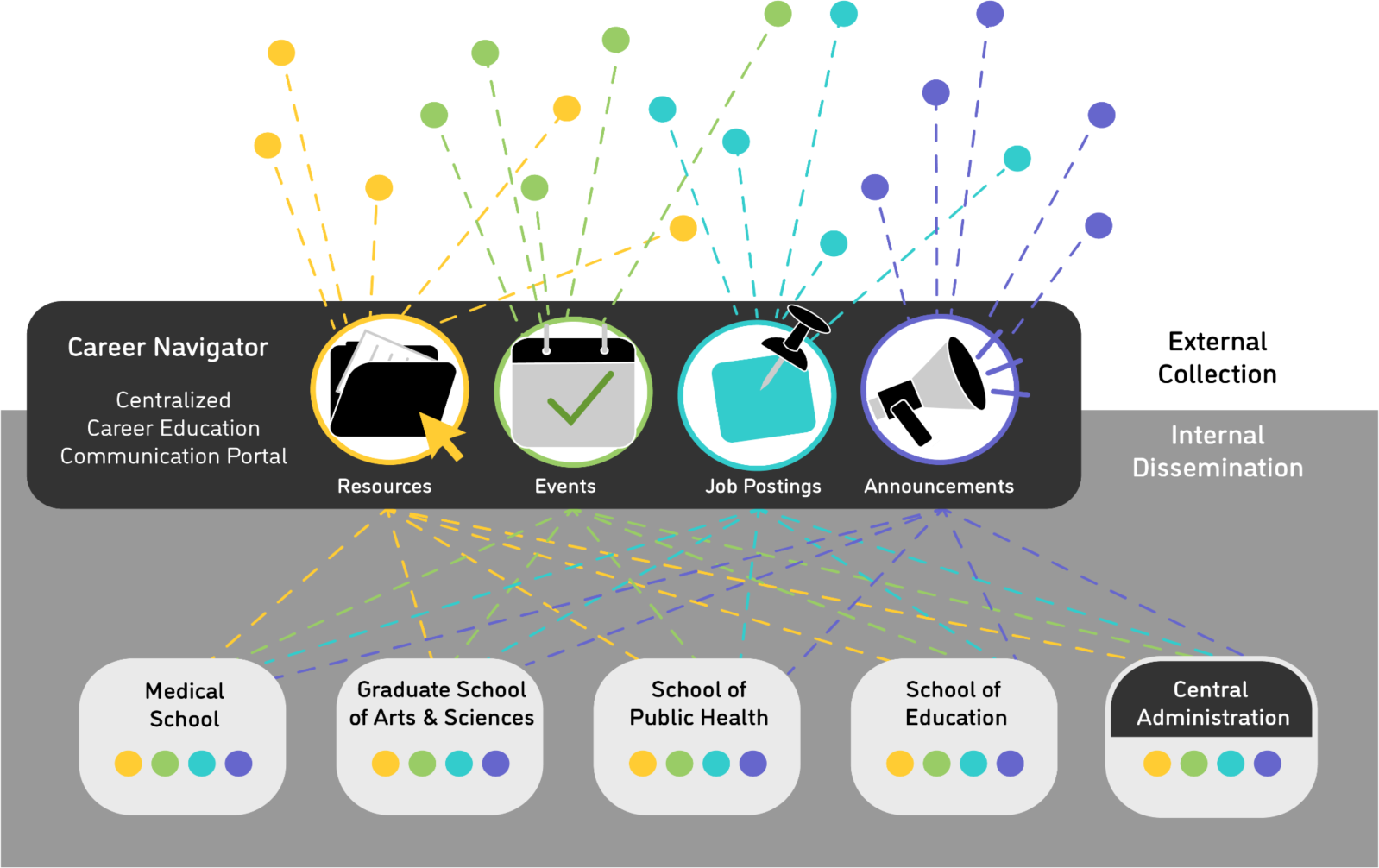
Model of integrating essential career education information to graduate students embedded in a complex landscape of providers and resources. This information includes announcements, resources, events and job postings. These can emerge from internal or external sources/providers. Internal providers may be any school within a university, and can reflect many different types of offices or organizations in each of these schools. In addition, some essential career education information and support may emerge from central administration such as a Provost’s office. A centralized career education communication portal allows for cross-school information sharing within an institution; additionally, it allows managers to centralize and organize information and resources originating from providers outside the university.

To collate resources (on the Resources page), we similarly applied “chunking” to the development of a taxonomy for resources. The taxonomy we developed for the Career Navigator is built around seven nodes (**Table 1**). These nodes are (1.) *Charting Your Course*, (2.) *Wellness and Safety*, (3.) *Finding and Securing Funding*, (4.) *Academic and Professional Skills*, (5.) *Interpersonal Skills*, (6.) *Career Exploration*, and (7.) *The Job Search*. To date, all the resources have been assigned to one or more of these nodes, which are further subdivided into subnodes. For example, the *Academic and Professional Skills* node includes subnodes for *Research Skills*, *Oral Communication* and *Business Skills,* among others. This infrastructure of nodes and subnodes allowed us to collate more than 300 resources into a well-organized database. These include guides, videos, and online courses. To better explain the taxonomy, we outline two representative, publicly available resources from each node (as presented in **Table 1**).

**Table 1:**
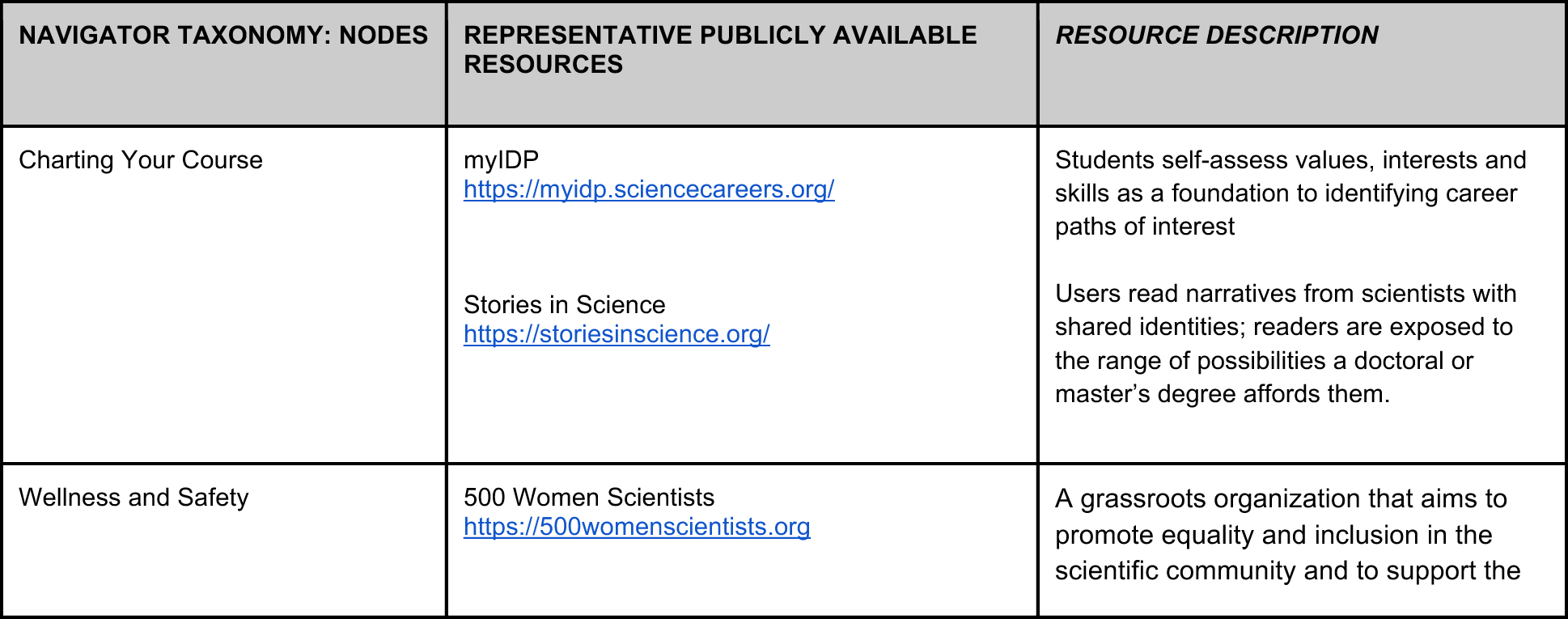
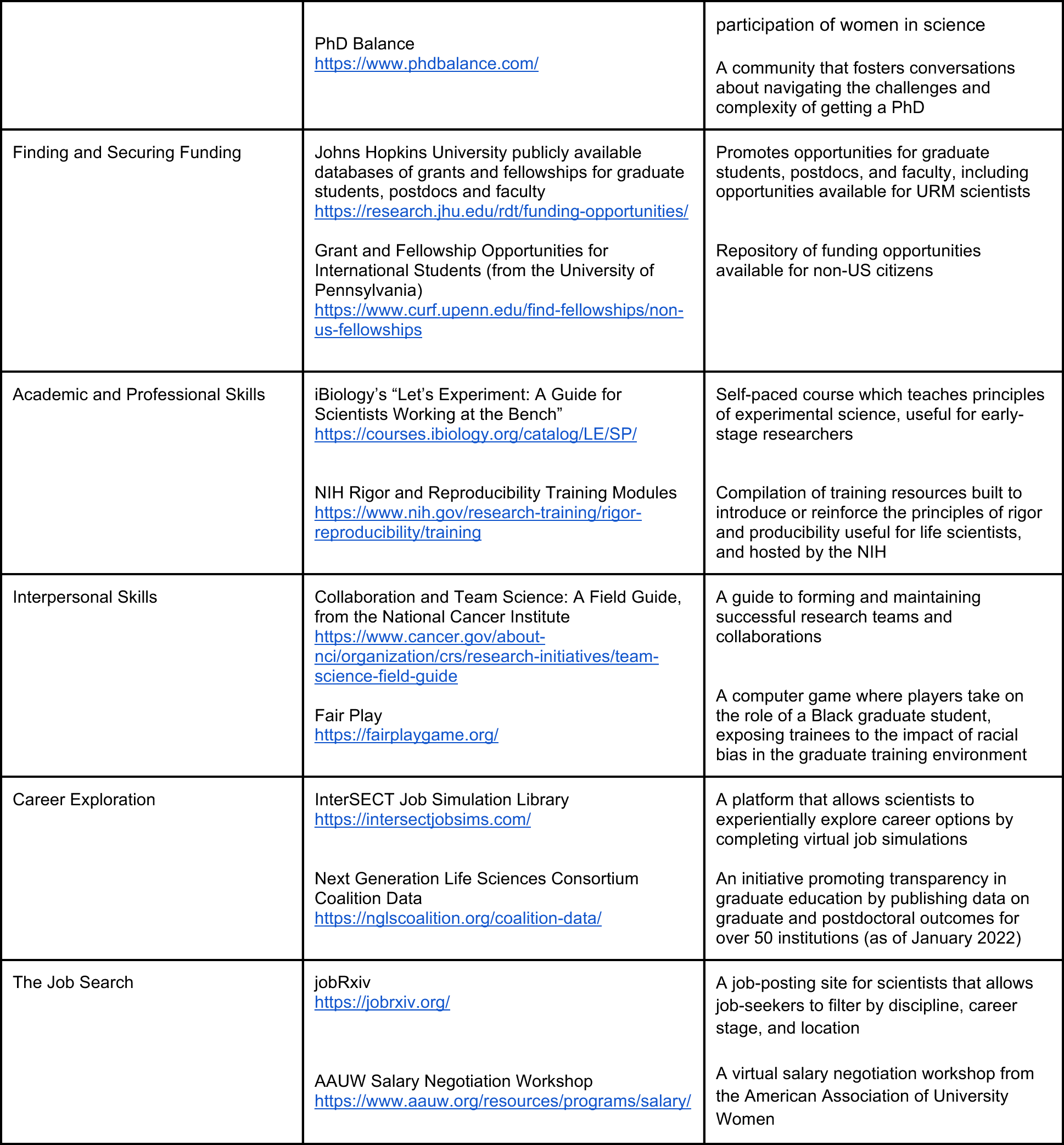
Career Navigator resource taxonomy, featuring its 7 major resource nodes, and highlighting two representative resources and for each node as collated on the site.

The *Charting Your Course* node encompasses resources trainees can use to broadly shape their graduate experience and identify their academic career direction. Under *Charting Your Course*, we feature *myIDP* [11] and *Stories in Science* [12], which both provide examples of, and guidance towards, shaping individual growth in professional development and career exploration.

The *Wellness and Safety* node contains resources that support student mental health and well-being, financial well-being, personal/public/lab safety, and inclusion and belonging among other themes. Here, we feature *500 Women Scientists* [13] and *PhDBalance* [14].

In the *Finding and Securing Funding* node, we feature the Johns Hopkins University funding opportunities website [15] and a webpage from the University of Pennsylvania that lists funding opportunities for non-US citizens [16], as well as notes on applying to various fellowships and grants.

The node *Academic and Professional Skills* includes resources that teach research, quantitative, written and verbal communication, and business skills. This includes a video series from iBiology, “Let’s Experiment: A guide for Students Working at the Bench” [17], and the NIH training modules designed to improve rigor and reproducibility in research [18].

In the *Interpersonal Skills* node, we include resources that support trainees in their soft skill development including diversity, inclusion, and avoiding bias; teamwork and collaboration; mentoring; and conflict resolution. Two exemplary resources from this node are “Collaboration and Team Science: A field guide” [19] and “Fair Play” [20].

Under *Career Exploration*, we include resources trainees can use to identify post-graduate career options, investigate their personal personal and professional values, explore various career options, and choose their next step. Among other resources, we feature the InterSECT job simulations library [21] and the data available through the Coalition for Next Generation Life Science [22].

*The Job Search* node contains resources for trainees actively seeking post-graduate employment. Two featured resources are jobRxiv, a platform with job postings targeting scientists, [23] and “Work Smart & Start Smart: Salary Negotiation” [24].

Building a comprehensive and organized website is a first step; however, students need reasons to continuously visit the site - for general utility and to engage with new content (resources, events, job postings, and announcements). Once the Career Navigator was launched at the start of the fall 2018 semester, we capitalized on new student orientation activities to introduce incoming students to this website. Additionally, we used emails from central administrative offices (in the graduate school and office for education at the medical school) to notify all active students of this new centralized portal. We communicated directly with graduate program directors and managers to encourage them to notify students about the platform. By attending faculty-student retreats and presenting on the platform, we also spread the word in-person and simultaneously to students, faculty and administrators. Career advisors who meet with master’s and doctoral students were also made aware of these platforms and encouraged to review with students during workshops and 1-on-1 meetings.

To promote ongoing use, we encourage visits to the site and related content through three avenues: themed events, an email digest, and social media. Next, we describe our intentional approaches and best practices in using these channels to encourage site engagement.

Exemplary events aligned with each node are presented as an organized “para-curriculum” over the course of the academic year (**Table 2**). A review of some exemplary events we have hosted - some in-person and some virtual - highlights how we revisit these themes and remind students of their importance in their professional development.

**Table 2:** The resource taxonomy, featuring its 7 major resource nodes, highlighting two representative theme-aligned events for each node as managed through the Career Navigator. Event advertisements make clear the nodes they represent and which run over the course of an academic year. (L = events we have run live/in-person, V = events we have run virtually).

| NAVIGATOR TAXONOMY: NODES | REPRESENTATIVE EVENTS | EVENT DESCRIPTION |
| --- | --- | --- |
| Charting Your Course | CV of Failures [L]<br>(related articles Stefan 2010, Johannes Haushofer CV of Failure 2016) | Event dedicated to discussions on the value of resilience along with real, personal experiences of overcoming failure in navigating one's career path |
|  | Finding and Choosing Mentors and PIs [L] | Event designed to support early-stage students as they complete rotations and make the critical choice of scientific research advisor, featuring senior graduate students serving as peer mentors in the structured discussion |
| Wellness and Safety | Ergonomics for the Lab and Workplace [L]<br>Overcoming Impostor Syndrome [V] | Introduces proactive strategies to maintain physical health, with a focus on workplace ergonomics; inspired by suggestion from a late-stage graduate student - via response to a standard annual program evaluation survey - as an early-stage intervention<br>Introduces and explains the concept of impostor syndrome as a foundation for discussing strategies for identifying and overcoming it toward success in graduate school |
| Finding and Securing Funding | Foundations of Grant Writing and Applying for Fellowships [L,V]<br>Strategies and Resources for Landing a Doctoral Fellowship (NSF, NIH T32, NIH T32D) [L,V] | Designed for both live and virtual delivery, the session provides attendees with an overview of the grant-writing process.<br>Intended for later stage graduate students, session is focused on securing a postdoc after graduation, with panelists sharing advice including strategies for applying for competitive postdoctoral fellowships and specifically designed to highlight postdoctoral experiences in different sectors (e.g., academia vs industry) |
| Academic and Professional Skills | R3 and Me: A Toolkit for Rigorous and Reproducible Research; Introduction to Electronic Lab Notebooks [V]<br>The Power of Persuasion: A Public Speaking Workshop [L,V] | A seminar touching on the importance of rigor and reproducibility in research and relaying resources for enhancing competencies in these areas<br>Lays emphasis on the communication skills necessary to deliver an effective presentation, incorporating live practice activities |
| Interpersonal Skills | How and Why to be a Good Graduate Peer Mentor [V]<br>More Than Just Paperwork: Making IDPs part of a Meaningful Mentor-Mentee Discussion [L,V]<br>(related article Vincent et al. 2015) | Interactive workshop covering the topic of peer mentoring and its positive impact on mentees; this workshop is now a foundational workshop for an extended series of workshops designed for graduate peer mentors<br>Emphasizes the importance of individual development plans (IDPs) as well as effective strategies for preparing, executing and following up on an IDP discussion |
| Career Exploration | Identifying Your Ph.D. Transferable Skills [V]<br>Backward Career Design: Using the Reality of Data and Outcomes to Plan Your Next Steps [L] | Teaches attendees how to effectively self-reflect and map the skills relevant to many different career paths that may be of interest<br>Workshop built around emerging data on graduate professional development and outcomes, including findings on the factors influencing choices in graduate school and from graduate bioscience alumni |
| The Job Search | Making a Case for Yourself: Crafting an Effective CV [L,V] | A workshop introducing students to the best practices in developing a curriculum vitae (CV) for successful job applications, contextualized as part of the academic job packet |
|  | Global Mobility for U.S. Trained Researchers: Challenges and Resources [V] | Offers advice on preparing for success outside of the U.S. research ecosystem, including highlighting the rewards and challenges of a career overseas and also in service of our international scholars who are considering paths to return to their home countries for their next professional steps |

Aligned with the theme *Charting Your Course*, several events and workshops have been presented to help trainees transition into and plan their development in graduate school including the “CV of Failures: A Scientist’s Guide to Resilience” and “Choosing Mentors and Mentoring Up.”

The events aligned with the *Wellness and Safety* node promote the overall well-being of graduate students; for example, the “Ergonomics for the Lab and Workplace” and “Overcoming Imposter Syndrome” workshops.

The *Finding and Securing Funding* related events are designed to best inform students of scholarships and fellowships during graduate school. These include workshops on the “Foundations of Grant Writing and Applying for Fellowships” and “Strategies and Resources for Landing a Doctoral Fellowship.”

The *Academic and Professional Skills* theme enables numerous events, including those geared towards developing research and communication skills. “R3 and Me: A Toolkit for Rigorous and Reproducible Research” and “The Power of Persuasion: A Public Speaking Workshop” are exemplary workshops.

Workshops related to the *Interpersonal Skills* node are aimed at sharpening soft skills essential for academic and professional development; for example, sessions on “How and Why to be a Good Graduate Peer Mentor” and “More Than Just Paperwork: Making IDPs part of a Meaningful Mentor-Mentee Discussion.”

For the *Career Exploration* theme, we host multiple workshops and events geared toward helping students identify their values, career options, and ultimately the right career track for them. These include “Identifying Your Ph.D. Transferable Skills’’ and “Backward Career Design: Using the Reality of Data and Outcomes to Plan Your Next Steps.’’

For the *Job Search* node, several seminars and workshops tailored toward the job application process are featured. This is illustrated through the sessions “Making a Case for Yourself: Crafting an Effective CV” and “Global Mobility for U.S. Trained Researchers: Challenges and Resources.”

Where possible, we pre-circulate relevant literature related to the events. For example, at the start of the “CV of Failures” event, we highlight not only the article that first proposed the value of faculty creating their own version of these “failure CVs” [25] but also one of the most famous examples of a faculty member who took up the challenge [26]. Likewise, in preparation for the discussion on effectively using IDPs as part of mentor-mentee discussions, we highlight a paper describing best practices [27].

These themed events were added to a landscape that already featured graduate professional development programming, and were designed to be complementary. We note, where possible in advertising, the alignment with these other existing programming and the themes in our framework. This includes the aforementioned series of workshops for graduate peer mentors, which reinforces the goals of our *Interpersonal Skills* node. Our (previously) credit-bearing PhD PATHFinder course is aligned with the *Career Exploration* node. Another series of workshops presented as *Preparing the Academic Job Packet* focuses on the *Job Search* for the academic science career path.

To keep students updated on changes to our site, we started a new email digest in April 2020 (coincidentally just after we transitioned to remote learning and work). Since then, we have sent out emails using an email management system (MailChimp). These bi-weekly messages (monthly during breaks) allow us to consolidate messaging to our students, summarizing information in an email (for which students and other users opt-in and can unsubscribe) but not compromising our goal of limiting the frequency of email. These digests have typically highlighted upcoming events (including direct registration links) as well as new resources we have posted to the site. Readers are encouraged to check for updated job postings and announcements. (We discontinued this version of digest in 2022 as we focused on upgrades to the site.)

We use one social media channel, Twitter (@HMSCareerNav), to connect with our students and other followers. We launched this Twitter account in February 2020. We use our Twitter account for frequent updates on new resources and articles added to our site. The Twitter account also allows us to highlight individual job postings and announcements (where relevant to the broad public audience). We also tweet final reminders about upcoming events (within 1-2 days) as a final push for our audience to consider registering for and attending our events.

Through the Contact Page on our site, users are directed to submit information for review and posting via our email address. The site was initially maintained by graduate students who collaborated on the build of the site, but as volume increased, it necessitated the hiring of a full-time staff member whose responsibilities include the maintenance of the site and coordination of other activities which help drive site use. These other activities include the aforementioned events and job postings management. The process of managing and posting this information necessitated and bolstered the value of a taxonomy for the collection of resources. Resources that are submitted to the site have to be organized into a useful framework to make search and navigation easy (reflected as the rows in **Table 1** and **Table 2**).

### Tracking engagement for feedback and optimization

Since launch, we have monitored engagement with our site using Google Analytics. To understand how engagement has changed over time and responded to our innovations, we analyzed the usage patterns over the initial three years - the period spanning July 1, 2018 to June 30, 2021 (**Figure 3**). Our analysis ends in June 2021 as the end of the last complete academic year before we began analysis. Encouragingly, we observed increasing engagement over time. The “Users” plot (**Figure 3A**) shows the number of users on the site for each day; it shows large fluctuations in our users which can be attributable to many things. For example, large spikes could coincide with days when we run workshops to teach users about the features on the site, wherein multiple people will simultaneously log to the website to explore. In contrast, the “Active Users” plot (**Figure 3B**) shows three traces with the number of users, over time, in the 1-day increment (bottom trace) which is the same as data presented in **Figure 3A**; the total number of users for the past 7 days for each date (middle trace); and, the total number of users for the past 30 days for each date (top trace). In **Figure 3B**, the orange arrow designates the start of the COVID-19 pandemic which also marks the start of a large increase in usership for our site. (Note that a single user who logs in multiple times per day from the same device is only counted as a single “user,” despite having many “sessions.”)

**Figure 3.**
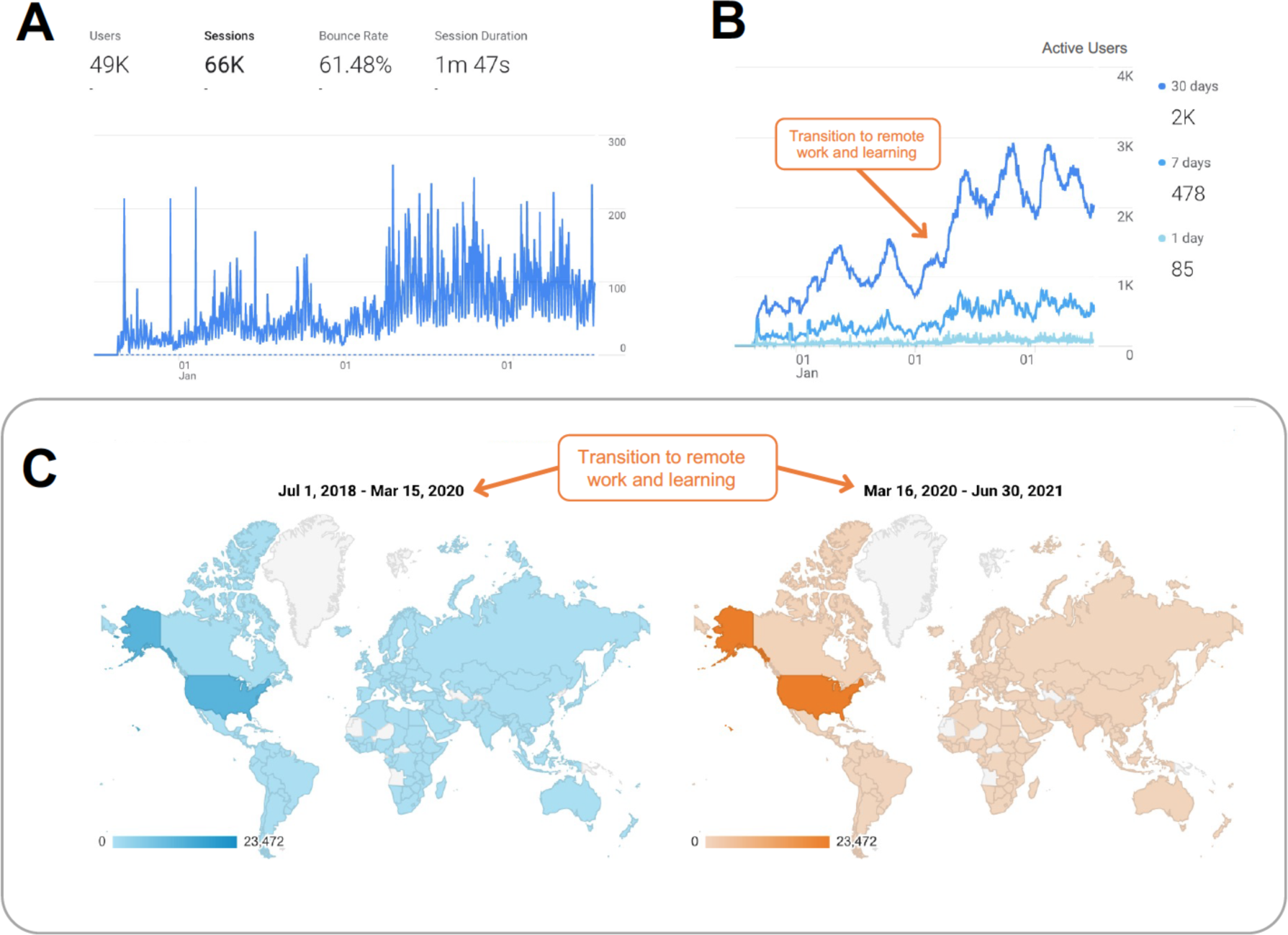
Career Navigator site use for the period spanning Jul 1, 2018 through June 30, 2021. **(A)** User interaction with the site varies widely from day-to-day. The peaks represent high activity on the site associated with event registrations and email or social-media based campaigns encouraging students to check out new resources, events, job postings and announcements on the site. Over a three-year period, ∼66,000 interactions were logged to the site. **(B)** Active User counts, the average users of the last 30-day period, can provide a more meaningful metric for the site given interactions from email digest and social media are not instantaneous: an email or Tweet may not be read for days after distribution though they may still drive site engagement. The orange arrow highlights the start of remote teaching and learning associated with the COVID-19 pandemic. **(C)** Geographic distribution of users on the site in two periods separated by the start of the COVID-19 pandemic as reflected at our R1 institution: July 1, 2018 - March 15, 2020 (pre-COVID-19) and March 16, 2020 - June 30, 2021 (COVID-19 pandemic). In both periods, the largest number of users were present in the United States but users were detected on all continents and from almost every country in the world. As such, the pivot to remote work and online learning (including our many students who remained in their home countries) did not explain the global usage of the site.

The users who were accessing our site were widely distributed across the globe, with engagement coming from almost every country in the world (**Figure 3C**). The distribution patterns before and after the start of the COVID-19 pandemic, which affected the geographic location of many of our users, appear similar. Contrary to our expectation that we would see broader geographical usage after the start of the pandemic, the two distribution patterns (before vs after March 15/16, 2020) appear relatively similar. In both cases, the largest number of users were located in the United States. It is not possible to use this data to evaluate fine-scale changes as the color densities at lower intensities are not distinguishable by eye. Therefore, we cannot draw conclusions about relative use in other countries. Those conclusions would also be confounded by the relative populations in those countries.

In order to better understand the impact of our iterative enhancements, we focused on “Active Users” across the 6-month increments making up our 3 years of data (**Figure 4**). First engagement with the site begins near the end of August 2018, as the site was launched as part of student orientation and the start of the fall semester (**Figure 4A**). In the period of January 1, 2019 - June 30, 2019, usage of the site began to increase noticeably around March 2019 when the first versions of the Professional Development Workshops (PDWs) were offered. Registration was managed on the site which likely explains the increase in users which began around this time and continued as the PDWs sustained attention to the site (**Figure 4B**). Usership of the site fell over the summer of 2019, due to both a lack of programming and likely a general reduction in the number of students accessing such websites over summer periods (addressed below). In fall 2019, starting in September, active user numbers began to rise again (**Figure 4C**) as professional development programming resumed, this time in the form of Professional Development Days (PDDs). The PDDs presented 2-3 PDWs of similar themes consecutively on one afternoon, which made it easier for students to block out the time and travel to campus if necessary.

**Figure 4.**
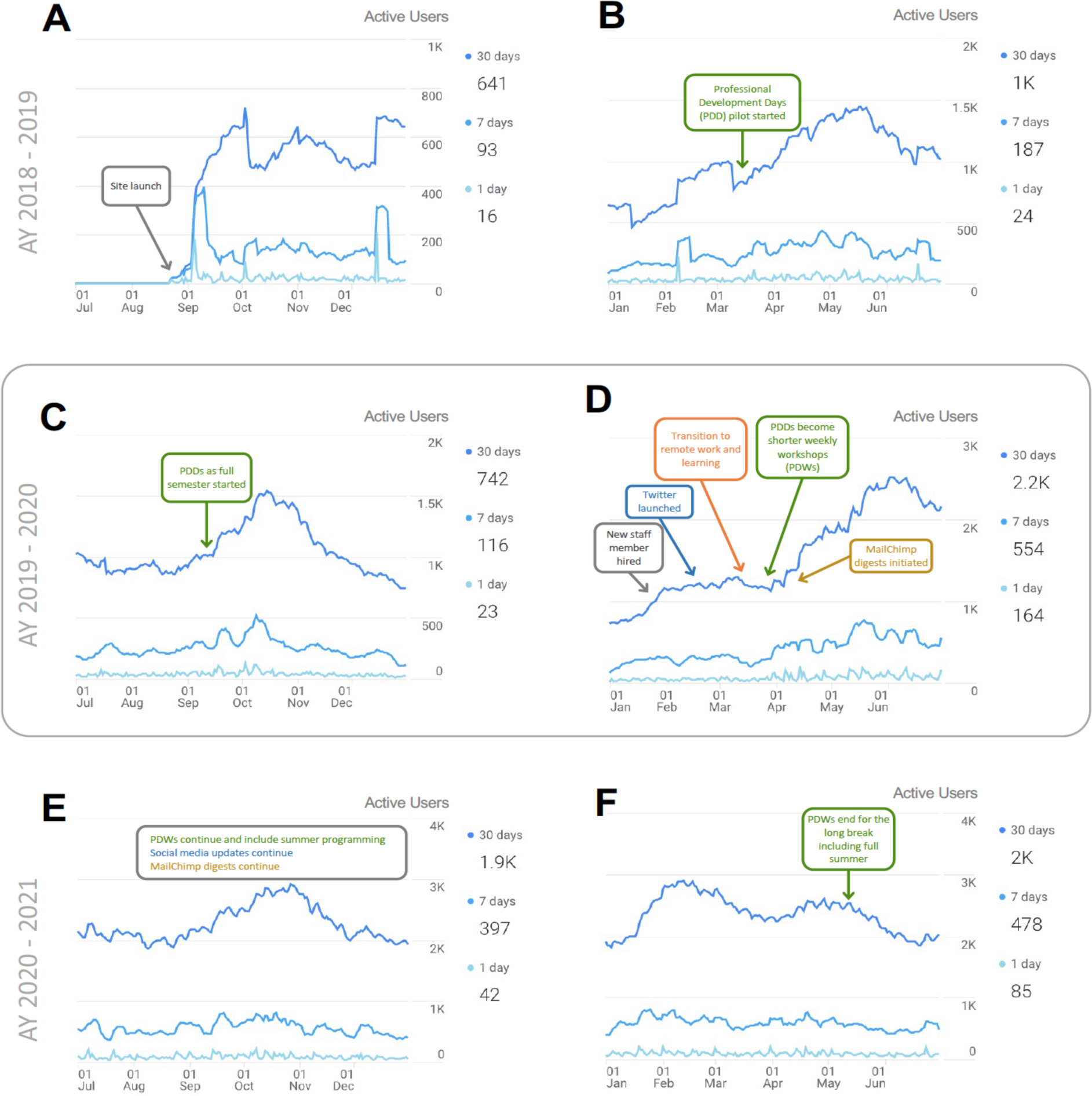
Career Navigator site Active Users in 6-month increments spanning Jul 1, 2018 through June 30, 2021. **(A)** Active users from July 1, 2018 through December 31, 2018. In late August 2018, the first activities in the site register as we conduct user testing and launch the site in alignment with new student orientation and the start of the fall semester. The site was also shared with all students via email and through presentations at departmental gatherings including students, faculty and staff. **(B)** Active users from January 1, 2019 through June 30, 2019. Site usage increased during this period, when the first two professional development days (PDDs) were piloted in-person for master’s students and three workshops were offered for doctoral students, with registration managed through the site. **(C)** Active users from July 1, 2019 through December 31, 2019. Highest usage on the site was in late September through October which coincided with weekly PDDs. Each PDD featured 2-3 consecutively executed workshops sharing similar themes. **(D)** Active users from January 1, 2020 through June 30, 2020. In order to scale the service and programming offered by the Career Navigator site, a new full-time staff member was hired (early February 2020) and this was a few weeks before the start of remote work associated with the COVID-19 pandemic (mid-March 2020). Only one PDW was held in-person before the transition to online professional development programming for the rest of the Spring. Online professional development programming increased usage of the site nearly two-fold following the pivot to accommodate remote learning and work. **(E)** Active users from July 1, 2020 to December 31, 2020. Usage of the site remained high with active user counts hovering between 2K and 3K users for most of that period. The site was featured during Orientation events for new students, which likely drove users to the site along with a small number of PDWs offered from September through December. **(F)** Active users from January 1, 2021 through June 30, 2021. Activity on the site was sustained between 2K and 3K users per 30-days for the duration of this period, to which Professional Development Workshops (PDWs) contributed.

One PDW was held in person in March 2020 before we transitioned to remote learning due to the COVID-19 pandemic. Thereafter, our programming was delivered as weekly workshops given the constraints of virtual engagement and the gradual rise in “Zoom fatigue” [28], a form of mental exhaustion and burnout experienced from prolonged and frequent use of video conferencing and virtual communication tools. Multiple other improvements were made during this period, January 2020 - June 2020 (**Figure 4D**): we hired a new full-time staff member who could support this site (February 2020), launched our Twitter account (February 2020) and started the new email digest summarizing upcoming events and new additions to the site (April 2020). In summer 2020, we also offered some - even though more infrequent - online programming, assuming that our audience would rather be taking advantage of the opportunity to be outside after months of restrictions to their homes amidst social distancing.

The usership of the site was maintained at elevated levels owing in part to continued virtual professional development programming over the periods July 2020 - December 2020 (**Figure 4E**) and January 2021 - June 2021 (**Figure 4F**). When active users are viewed in aggregate, it is clear that while usership is sustained at high levels, there are dips close to the end of June and end of December each year, other than for June 2020 when users were actively engaging in online work due to the pandemic (**Figure 3B**). This lower activity on the site is likely related to our reduction in programming and announcements, as student and staff activity are reduced during the breaks.

Although the next academic year was not complete at the time of analysis, the usage patterns have shown sustained user engagement of >2,000 users per month for the academic years 2021-2022 and so far in 2022-2023 (data not shown). This is despite a reduction in our programming and email digests during the 2021-2022 academic year. This is also true so far for the 2022-2023 academic year while our programming has increased again but our email digests have not been distributed as we adapt to new integrations on the Career Navigator. For the purposes of this analysis, we measured “engagement” as the number of interactions with the website as a whole (i.e. counting the number of times the website was visited in total). This was in-line with a high-level study of the site, although we recognize that there are interesting questions to address in the future such as the patterns driving repeat visits for users, the time spent on the site (and which pages tend to retain users longer), and which sub-pages, including pages for each resource, are visited most frequently. This includes analysis of metrics such as “bounce rate” and “time spent on the site” which are visible in **Figure 3A**. Similarly, Google Analytics data shows that among our top 10 page views in the same period as the data presented above, the most frequently visited page on our website (not surprisingly) is the homepage, and likewise that each of the main subpages (Resources, Events, Job Postings, Announcements, and Contact Us) are within the top 10 (data not shown). The other 4 pages making up the top 10 are the *Career Exploration* page (#10) from our *Resource* taxonomy, and three specific resource pages: the Graduate School of Arts & Sciences Business Club Mini-MBA page (#2), the Health Professions Recruitment and Exposure Program (#7), and the Journal of Emerging Investigators (#9).

## DISCUSSION

### Summary and context of data vis-à-vis research questions

Doctoral students across all years of our programs expressed a preference for centralizing professional development and career education information online - showing a higher preference for this mode of communication over digest emails or separate emails for each notice, whether at the office/institution or program level (**Figure 1**). Prior to data analysis, we assumed that the preference for a single website would be observable (or most pronounced) in more senior students (e.g., fourth-, fifth- and more senior-year students) given the time accumulated receiving multiple channels of communication; however, we found that even second year and third year (early-stage) doctoral bioscience students shared the perception of this need (**Figure 1**). The launch of the Career Navigator (August 2018) demonstrated our capacity to translate our model for a consolidated career education portal into a real, accessible, student-friendly website (**Figure 2**). The site is publicly available and usage patterns show steady and sustained increase (**Figure 3B** and **Figure 4**). Instantaneous (single day) measures of activity can be useful to track interventions with large audiences; for example, in demo sessions (**Figure 3A**). While we could not use geographic distribution to answer questions about changes in usage pre- vs post-COVID-19 pandemic, the overall geographic distribution highlights wide access and use of the site (**Figure 3C**).

A key aspect of the site was the development of a 7-part taxonomy to organize resources (**Table 1**), which in turn served as a foundation for designing our para-curriculum of professional development programming (**Table 2**). The events have evolved over time: we have transitioned from stand-alone in-person/live workshops, to single-day multiple-workshop in-person sessions, and then to stand-alone virtual sessions. We recently began offering in-person workshops again (September 2022), though the majority of our workshops remain virtual. This programming and other interventions - the hiring of staff to support the site, new communication channels such as an email marketing service and Twitter - coincided with increasing use of the site which is most dramatically observable around March 2020 (**Figure 3D**). From May 2020 through May 2021, the total users within any given 30-day period have been sustained above 2,000 users (**Figure 3B** and **Figure 4D-F**). We have observed sustained user engagement >2,000 over sequential 30-day periods for the 2021-2022 academic year as well and into this 2022-2023 academic year so far.

### Limitations and Future Directions

To address our research questions, we used program evaluation data to show that our students prefer a centralized career education site to track and find information relevant to their professional development. We acted on the need for a centralized portal, which led to the development of the Career Navigator, implementing best practices we learned during the due diligence process and partnering with an expert in web development as a consultant. A primary outcome of the building of this site is the development of a new taxonomy to organize resources along seven nodes and their subnodes.

Since launch, we have used multiple touchpoints including orientation events and faculty-student retreats - in addition to traditional email blasts - to reach students, faculty and staff who were the stakeholders for this website. Going forward, we are planning on using additional channels to drive further engagement on the site, such as by encouraging our graduate programs to include a brief description of the website and its link on all IDP forms that doctoral students complete annually. Likewise, we can encourage the inclusion of similar information on Dissertation Advisory Committee (DAC) forms, which doctoral students complete annually when they meet to discuss research progress and plans. Similar and multiple other strategies to enhance partnerships between research faculty and career educators have been recently described [29]. Moving forward, we plan to use program evaluation data to assess whether the Career Navigator is used during IDP discussions and makes finding information more *efficient* for graduate students and mentors.

Efforts to continuously draw users to the site have been successful, including events themed to align with the taxonomy, event registration, bi-weekly email digests on updates and the coupled social media.

While we can use Google Analytics to determine what pages and subpages have been most frequently viewed, we do acknowledge some limitations. For example, while we can identify the resources most viewed on the site (e.g. the Business Club Mini-MBA program), there are confounding factors that need to be considered in using this information. For example, specific subpages on our site can be pinned elsewhere; the Business Club Mini-MBA program page can be linked to blogs for those interested in business school, which drive up the views on this specific page. A new platform, Career Cognitive City (described below), removes some of these biases and may better address questions on the resources that are searched for and viewed by users navigating the site. Similarly, in line with policies at our institution, we have been highly selective in the addition of wellness and safety resources to the platform, which influences any interpretation of the relative use of mental health and wellness resources embedded in a professional development matrix.

### Generalizability, Transferability and Broader Implications

The scope of career guidance and preparation for life science graduate students has broadened. There is now widespread recognition from graduate program directors, faculty advisors, students, career services staff, and funding agencies that students should be encouraged - early and throughout their training - to explore the range of professional opportunities and identify their transferable skills. Career advisors and coaches are now called upon to not only deliver training, create resources and run events aimed at career exploration, networking and the job search, but to also cover topics and trainings on wellness, work-life balance, grant writing, general communication, growing mentor networks, and culturally aware job searching. Graduate career education, then, traverses the full graduate student experience and requires stage-specific resources to aid in a broad range of training. To maximally benefit the student, these resources need to be available on-demand, organized, and easily accessible for students when they are seeking them. Here we have described an approach to addressing this challenge by building the Career Navigator, which collates these on a central website. We also describe the taxonomy we developed to help students effectively search and find the resources they are seeking.

The approach we have taken to organize graduate career education information resonates with the call to accessibly organize mental health resources [7]. Moreover, as noted above, providing support for trainees’ well-being and mental health - as well as teaching them approaches to maintaining these for themselves - have also become integrated into career education. For example, we commonly discuss the need for training students on resilience and overcoming failures; these discussions complement ongoing general discussions pertaining to mental health and navigating career hurdles and setbacks.

The relationship between mental health challenges graduate students face and career preparation, in both the traditional and the broader scopes presented here, has been documented. Graduate students’ perspectives on their career prospects are strong predictors of their sense of satisfaction and incidence of depression [6]. Helping trainees and faculty mentors build and sustain healthier mentoring relationships; build supportive communities to support graduate students who identify as transgender, non-binary or women; improve their approach to work to better manage work-life balance; and better support under-represented students, all translate to better mental health for graduate students [30]. Likewise, these interventions align with improved career outcomes for graduate students. Related to this mission, it is important to emphasize to faculty the research findings that show that metrics such as time-to-degree, total publication number, or first author publications are not negatively affected by participation in professional development activities [31].

A related consideration is that of information overload: as more and more resources become available, and our graduate career advising community works to spread the word about these resources through traditional modes of communication (e.g., email), are we contributing to information overload and inadvertently having a negative impact on students? This communication overload may negatively impact student well-being and mental health, or discourage students from meaningful engagement with career planning if it is perceived as overwhelming. We could not find published studies that investigate this phenomenon in graduate students. However, there has been ample consideration of information (or digital) overload in the workplace [32] including how it affects focus and attention to work [33] and how employee engagement with email communication can be a predictor of workplace burnout [34].

The career and professional development website described here is an institutional response to efficiently and effectively collate professional development information for bioscience graduate students. Websites can be useful places to aggregate information regarding mental health resources, but their impact is amplified through mechanisms that encourage graduate students to visit the site frequently [7]. Considering the relationship between graduate student mental health and professional development, the same principles should be relevant to websites built to promote graduate student professional development. We utilized a combination of approaches to encourage return visits to our site including frequent content updates such as announcements and event registration as well as linking back to the site via social media (Twitter) and in bi-weekly email digest powered by an email marketing service (Mailchimp). Importantly, we committed resources to manage these interventions through the hiring of a full-time staff member whose work includes managing the Career Navigator, vetting inquiries, managing logistics for programming, and handling relevant communication.

Our experience with the site thus far has led us to consider ways to enhance the student experience with this platform. We are now (since September 2022) embarking on a “next-generation” version of this site, incorporating technological integrations that enhance the user experience with each page on our platform. For example, our *Resources* page started off with ∼90 resources spread across its 7 nodes; as of January 2022, there were more than 300 resources on the site. We realized the value of a more dynamic visual interface to present these resources to students and to take advantage of advanced cataloging features to help make them customizable and searchable by different target audiences and their needs. Given the resource investments in building this repository, it was important to make such a library accessible to any trainee of interest - including those outside our institution - particularly as we also highlight so many excellent resources from other institutions. We developed a new platform for this purpose, called Career Cognitive City (abbreviated Career CogCity) which we recently launched [35]. Career CogCity is a free-standing public platform, which for the convenience of our students has been embedded in our *Resources* page as well.

As another example of enhancing a page on our site, we are capitalizing on the functionality of a Career Services Management (CSM) platform to expand our *Job Postings* page so it serves as a better repository of internship opportunities and to allow for direct engagement between employers/recruiters and our students. We are also testing the relevance of the CSM to help us manage Individual Development Planning for students.

One of the unintended but important outcomes from the development of the Career Navigator is its taxonomy for resources. We have made many changes to our taxonomy since its launch. As we continue to collect resources, we import these and iteratively consider the validity and robustness of the nodes and subnodes defining the major organizational framework for this new platform. This process, in turn, informs our efforts to develop a *taxonomy guide for resources* similar to the one created for graduate bioscience career outcomes [36]. This taxonomy, which will continue to evolve as we validate, is reflected on our derivative platform Career Cognitive City.

Our motivation in this project, in related projects, and in dissemination here are three-fold: (a) student development, (b) faculty/mentor development and (c) staff and institutional capacity building.

With respect to **student development**, our goals have been to empower students to use the wealth of resources being developed to promote their professional development while doing so in ways that attenuate the possible impacts of digital overload through excessive emails. We see this resource library as a complement to individual development plans (IDPs), wherein this library of resources can be consulted at various stages of graduate school to harness “just-in-time” interventions that can help trainees overcome hurdles such as targeted skill development, finding relevant events across institutional boundaries, engaging with alumni, engaging with employers and finding jobs. We plan to continue to collect data on the use of this site to support our students’ engagement with IDPs.

By extension, we also see this as a vehicle for **faculty development**. One of the keys to graduate student mentorship should be a mentor’s prioritization of the trainee’s professional development. In some cases, mentors are wholly uncommitted to this goal. In other cases, mentors are keen to support trainees but do not have a top-level view of the many innovations being developed by the graduate advising community (across many institutions) or the breadth of knowledge about all the events being made available to trainees at their institutions (especially beyond the silos that often emerge at the departmental, programmatic or school levels). We hope that by lowering the entry barrier to engaging with resources, faculty will be more readily able to use them as part of their advising practice. By making both students and faculty mentors better informed, our goals are to enhance the mentor-mentee discussions that depend on IDPs.

Finally, by also sharing our approach for improving communication with our bioscience graduate students as well as our initial taxonomy for resources, our ultimate hope is that it may aid **staff development and institutional capacity building** at other institutions. The implementation of the new website was a low-cost, but high-time effort. Investments in these efforts have proven fruitful for us, given our priorities to support students in the overlapping issues of mental health, well-being, career education and professional development. We believe that educators at other institutions will find it similarly useful in applying our approach as a template for better supporting the needs of bioscience and other graduate students.

## METHODS

### Needs Assessment

To measure baseline interest in the development of a centralized career education platform for our graduate students, we took advantage of existing program evaluation data. We routinely collect survey data from our doctoral bioscience students (as we are establishing an annual survey protocol to gather continuous feedback as part of program evaluation).

We collected data from all second-year and above doctoral students (G2+ students) actively enrolled in our life sciences graduate programs at the end of the 2017-2018 academic year. Responses were collected from April 2018 through June 2018. This was a wide-ranging program evaluation survey, which included one question probing student preferences on career education communication: *“Please rate how effective each of the following methods is or would be to you, as a means of making you aware of resources concerning fellowships, career exploration events, internships, and job opportunities.”* Respondents rated the following suggested approaches using a 5-point Likert scale (not at all effective, slightly effective, etc.):

- *A centralized website hosting information [on] all regular and upcoming opportunities for life sciences students*
- *Separate emails from an administrative office (e.g., from DMS, HILS or PGE)* for each opportunity*
- *Separate emails from your dept* or program for each opportunity*
- *Weekly digest email from an administrative office (e.g., from DMS, HILS or PGE)* with all upcoming opportunities*
- *Weekly digest email from your dept or program with all upcoming opportunities*

(*Abbreviations: DMS = Division of Medical Sciences, HILS = Harvard Integrated Life Sciences, PGE = Program in Graduate Education, dept = department)

Our survey was distributed to all G2+ students using Qualtrics software. Student responses were incentivized by allowing students who completed the entire survey to opt into a raffle for one of many $50.00 gift cards. Our target population at the time was ∼1000 - 1100 G2+ students. Data were analyzed only for respondents who completed the entire survey (n = 329); this minimized the likelihood of having a single student respond more than once as in the example of a student who started the survey, allowed it to expire before completing, then re-started and completed the survey. (A % response rate for this data collection is misleading, given that the data collection period spanned April to June of the academic year when many of our G5+ students had or were defending their theses, and would have been very likely not to engage or to feel less motivation to provide feedback. It may also have coincided with other milestones such as Pre-Qualifying Exams which would be common for our G2 students).

### Website Design, Build and Aligned Programming

Two graduate students (co-first authors on this paper) were hired and included in all aspects of the website design to ensure that student perspectives were prioritized. The website was built using the OpenScholar platform, which was developed at our institution but is now commercially available. We were also able to procure a web development consultant from our University Office of Web Publishing, who worked with the design team on overall architecture of our site and the development of our inaugural resource taxonomy.

An informal focus group, dependent on convenience sampling for participants, was conducted prior to launch in August 2018. At that time, a small group of our faculty (∼6), mostly graduate program directors (e.g. Biological and Biomedical Sciences; Program in Neuroscience; Systems, Synthetic and Quantitative Biology) were provided with the link to the site and asked to provide feedback which was incorporated prior to launch.

Topics for our workshops - Professional Development Days (PDDs), Professional Development Workshops (PDWs) and Career Navigator Workshops (CNWs) as they have been called at various times - have been decided upon on an *ad hoc* basis including student suggestions and the identification of speakers through professional development networks (e.g., the Graduate Career Consortium [GCC]) or vendor solicitations.

For all in-person events, participant registration was managed through the native OpenScholar registration tool which collects basic information such as participant name and email address but is not customizable. Virtual events were delivered through Zoom. Registration for virtual events was managed using Google forms, which allowed us to collect additional questions as part of vetting individuals who signed up for our publicly accessible workshops while limiting threats such as “Zoom-bombing.”

In order to register for events, students have to visit the Career Navigator website; all announcements link to registration pages on the Events pages of the site rather than, for example, routing directly to a Google Forms registration page that would not cause students to visit the Events page. By ensuring students register on a relevant Career Navigator events page, Google Analytics can be used to measure the number of engagements with that page. It also increases the likelihood of off-target effects where students may see other upcoming events and consider attending those. Finally, for all events, advertisements are tagged with the relevant “theme” which is a node from the Resources taxonomy.

### Post-Launch Evaluation

We used the free Google Analytics tool to measure interactions with our site. A Google Analytics account linked to the URL for our site was established prior to launch in August 2018. Combined with the features of the OpenScholar platform, this allowed us to measure engagement with announcements, resources, events and registration pages, and job postings. The data we used in this analysis was aggregate data, and did not include information on individual users or their IP addresses. We have not utilized follow-up surveys, focus groups or interviews to measure student perceptions of the site. The survey data used in this study, which were collected as part of routine evaluation efforts, did not qualify as human subjects research as defined by our institution; hence, we do not have an IRB number to report.

For the period discussed here (July 2018 - June 2021), we did not conduct training (or learning) evaluations for our workshops due to limitations in personnel and logistical challenges in delivering evaluations to the right target audience.

## Funding

All funding for this project was institutional, including from the Harvard Initiative for Learning and Teaching (HILT) and the Office for Graduate Education at Harvard Medical School.

## Website Build

Gabriel Caro (Harvard University Information Technology) for collaborating on the design, development and implementation of the original Career Navigator, and for insights on the use and interpretation of Google Analytics.

## Feedback for Improvements

Countless Harvard graduate students, via pre-launch focus groups, surveys and in direct communication.

## Graphics

Olivia Foster Rhoades for graphic design (Figure 2) and visual editing (Figures 3 and 4).

## Pre-submission manuscript review and feedback

Thi Nguyen (the Science Communication Lab) and Rosalind Segal (Dean for Graduate Education at Harvard Medical School) for critical analysis and constructive feedback which enhanced this manuscript.

## Notes

### Competing Interest Statement

The authors have declared no competing interest.

## REFERENCES

1. Biomedical Research Workforce Working Group Report. 2012. National Institutes of Health. https://biomedicalresearchworkforce.nih.gov/docs/Biomedical_research_wgreport.pdf. [Last accessed December 19, 2023]

2. J. A. Hobin, C. N. Fuhrmann, B. Lindstaedt, P. S. Clifford, You need a game plan. Science (2012), doi:10.1126/science.caredit.a1200100.

3. R. N. Lenzi, S. J. Korn, M. Wallace, N. L. Desmond, P. A. Labosky, The NIH “BEST” programs: Institutional programs, the program evaluation, and early data. FASEB J. 34, 3570–3582 (2020).

4. National Institute of General Medical Sciences Predoctoral Institutional Research Training Grant guide. https://grants.nih.gov/grants/guide/pa-files/PAR-20-213.html [Last accessed October December 19, 2023]

5. National Institutes of Health Office of Intramural Training and Education website. https://www.training.nih.gov/ [Last accessed December 19, 2023]

6. The Graduate Student Assembly and the University of California-Berkeley, “Graduate Student Happiness & Well-Being Report 2014”, 2014 https://gradresources.org/wp-content/uploads/2015/09/wellbeingreport_2014-17.pdf [Last accessed December 22, 2023]

7. L. A. Krause, S. L. Harris, Get online to support wellbeing of graduate students. Elife. 8 (2019), doi:10.7554/eLife.53178.

8. J. Posselt, Promoting graduate student well-being: Cultural, organizational, and environmental factors in the academy. Council of Graduate Schools (2021) (available at https://legacy.cgsnet.org/ckfinder/userfiles/files/CGS_Well-being%20ConsultPaper%20Posselt.pdf).

9. J. Sablan, B. Mahoney, Reframing the Individual Development Plan. Inside Higher Ed, 2022 https://www.insidehighered.com/advice/2022/01/03/tool-toward-equity-graduate-student-career-development-opinion [Last accessed December 19, 2023]

10. The President and Fellows of Harvard College, Career and Professional Development Navigator. Hosted at Harvard Medical School, 2022. https://careernavigator.gradeducation.hms.harvard.edu/ [Last accessed December 19, 2023]

11. myIDP Science Careers Individual Development Plan. https://myidp.sciencecareers.org/. [Last accessed December 19, 2023]

12. Stories in Science. https://storiesinscience.org/ [Last accessed December 19, 2023]

13. 500 Women Scientists. https://500womenscientists.org [Last accessed December 19, 2023]

14. PhD Balance. https://www.phdbalance.com/ [Last accessed December 19, 2023]

15. The Johns Hopkins University. “Funding Opportunities” webpage. https://research.jhu.edu/rdt/funding-opportunities/ [Last accessed December 19, 2023]

16. The University of Pennsylvania. “Grant and Fellowship Opportunities for International Students” webpage. https://www.curf.upenn.edu/find-fellowships/non-us-fellowships [Last accessed December 19, 2023]

17. iBiology. “Let’s Experiment: A Guide for Scientists Working at the Bench”. https://courses.ibiology.org/catalog/LE/SP/ [Last accessed December 19, 2023]

18. The National Institutes of Health. “Rigor and Reproducibility Training Modules” webpage. https://www.nih.gov/research-training/rigor-reproducibility/training [Last accessed December 19, 2023]

19. The National Cancer Institute. “Collaboration and Team Science: A Field Guide”. https://www.cancer.gov/about-nci/organization/crs/research-initiatives/team-science-field-guide/collaboration-team-science-guide.pdf [Last accessed December 19, 2023]

20. The Fair Play Project and the Board of Regents of the University of Wisconsin. “Fair Play Game”. https://fairplaygame.org/ [Last accessed December 19, 2023]

21. InterSECT Job Sims and Contributing Authors. “InterSECT Job Simulations: Interactive Simulation Exercises for Job Transitions” webpage. https://intersectjobsims.com/ [Last accessed December 19, 2023]

22. Next Generation Life Sciences Consortium. “Next Generation Life Sciences Consortium Coalition Data” webpage. https://nglscoalition.org/coalition-data/ [Last accessed December 19, 2023]

23. jobRxiv webpage. https://jobrxiv.org/ [Last accessed December 19, 2023]

24. American Association of University Women. “Work Smart & Start Smart” workshops online. https://www.aauw.org/resources/programs/salary/ [Last accessed December 19, 2023]

25. M. Stefan, A CV of failures. Nature. 468, 467–467 (2010).

26. S. Haushofer, Johannes Haushofer: CV of failures (2016).

27. B. J. Vincent, C. Scholes, M. V. Staller, Z. Wunderlich, J. Estrada, J. Park, M. D. J. Bragdon, F. Lopez Rivera, K. M. Biette, A. H. DePace, Yearly planning meetings: individualized development plans aren’t just more paperwork. Mol. Cell. 58, 718–721 (2015).

28. R. Nadler, Understanding “Zoom fatigue”: Theorizing spatial dynamics as third skins in computer-mediated communication. Computers and Composition. 58, 102613 (2020).

29. S. Subramanian, J. A. Hutchins, N. Lundsteen, Bridging the gap: increasing collaboration between research mentors and career development educators for PhD and postdoctoral training success. Mol. Biol. Cell. 33 (2022), doi:10.1091/mbc.E21-07-0350.

30. T. M. Evans, L. Bira, J. B. Gastelum, L. T. Weiss, N. L. Vanderford, Evidence for a mental health crisis in graduate education. Nat. Biotechnol. 36, 282–284 (2018).

31. P. D. Brandt, S. Sturzenegger Varvayanis, T. Baas, A. F. Bolgioni, J. Alder, K. A. Petrie, I. Dominguez, A. M. Brown, C. A. Stayart, H. Singh, A. Van Wart, C. S. Chow, A. Mathur, B. M. Schreiber, D. A. Fruman, B. Bowden, C. A. Wiesen, Y. M. Golightly, C. E. Holmquist, D. Arneman, J. D. Hall, L. E. Hyman, K. L. Gould, R. Chalkley, P. J. Brennwald, R. L. Layton, A cross-institutional analysis of the effects of broadening trainee professional development on research productivity. PLoS Biol. 19, e3000956 (2021).

32. P. G. Roetzel, Information Overload in the Information Age: A Review of the Literature from Business Administration, Business Psychology, and Related Disciplines with a Bibliometric Approach and Framework Development (SSRN, 2020).

33. D. Dean, C. Webb, Recovering from information overload, (available at http://dln.jaipuria.ac.in:8080/jspui/bitstream/123456789/2179/1/Recovering%20from%20information%20overload.pdf).

34. C. P. Estévez-Mujica, E. Quintane, E-mail communication patterns and job burnout. PLoS One. 13, e0193966 (2018).

35. The President and Fellows of Harvard College, Career Cognitive City. Hosted at Harvard Medical School, 2022 https://hms-gradbiocareer.cognitive.city/public/resource-map [Last accessed: December 19, 2023]

36. C.A. Stayart, P.D. Brandt, A.M. Brown, T. Dahl, R.L. Layton, K.A. Petrie, E.N. Flores-Kim, C.G. Peña, C.N. Fuhrmann and G.C. Monsalve,. Apply inter-rater reliability to improve consistency in classifying PhD career outcomes. F1000Research 9(8), 2020. doi: 10.12688/f1000research.21046.2

